# SCA4 locus-associated gene Ronin (Thap11) increases Ataxin-1 protein levels and induces cerebellar degeneration in a mouse model of ataxia

**DOI:** 10.1101/2020.03.04.977405

**Authors:** Thomas P. Zwaka, Ronald Richman, Marion Dejosez

**Affiliations:** Department for Cell, Regenerative and Developmental Biology, New York, NY, USA; Black Family Stem Cell Institute, Icahn School of Medicine at Mount Sinai, New York, NY, USA; Department of Molecular and Human Genetics, Baylor College of Medicine, Houston, TX, USA; Jan and Dan Duncan Neurological Research Institute at Texas Children’s Hospital, Houston, TX, USA; Howard Hughes Medical Institute, Chevy Chase, MD, USA

**Keywords:** Ronin/Thap11, ataxia, Atxn1, SCA4, polyglutamine disease, Purkinje cells, embryonic stem cells, copy number variation

## Abstract

Spinocerebellar ataxias (SCAs) are a group of genetically heterogeneous inherited neurodegenerative disorders characterized by progressive ataxia and cerebellar degeneration. Here, we tested if Ronin (Thap11), a polyglutamine-containing protein encoded in a region on human chromosome 16q22.1 that has been genetically linked to SCA4, can be connected with SCA disease in a mouse model. We report that transgenic expression of *Ronin* in mouse cerebellar Purkinje cells leads not only histopathologically to detrimental loss of Purkinje cells but also phenotypically to the development of severe ataxia as early as 10-12 weeks after birth. Mechanistically, we find that Ronin is part of a protein complex in the cerebellum that is distinct from the one previously found in embryonic stem cells. Importantly, ectopically expressed *Ronin* raises the protein level of Ataxin-1 (Atxn1), the causative gene of the most common type of SCA, SCA1. Hence, our data provide evidence for a link between Ronin and SCAs, and also suggest that Ronin may be involved in the development of other neurodegenerative diseases.

## INTRODUCTION

Inherited ataxias are a group of autosomal dominant inherited neurodegenerative diseases characterized by loss of motor coordination and balance. Despite the genetic heterogeneity many ataxias show striking similarity and it is often difficult to distinguish between Spinocerebellar Ataxias (SCAs) based solely on clinical and pathological observations. The most common neuropathological feature of SCAs is the atrophy of the cerebellum that is accompanied by a characteristic loss of Purkinje cells. SCA1, at least five other SCAs (SCA2, 3, 6, 7 and 17) and other neurodegenerative disorders such as Huntington Disease (HD), are caused by proteins with abnormally long polyglutamine tracts as common cause in pathogenesis (Durr et al., 2010; Schöls et al, 2004; Paulson et al., 2009; Orr et al., 2007). Although, the origin of many other SCAs is still uncertain, there is an interesting polyglutamine repeat-encoding gene, Ronin (Thap11), located in a genomic region on chromosome 16q22.1 (chr16: 66,095,251 and 74,034,030; hg38) that has been mapped to SCA4 disease (Flaningan et al., 1996; Hellenbroich et al., 2003; Hellenbroich et al., 2006). Ronin, a transcriptional regulator that is essential for embryonic development (Dejosez et al., 2008; Fujita et al., 2017), seems worth to be considered in the context of ataxias for a number of reasons. First, consisting of 28/29 glutamine residues on average, it has one of the longest polyglutamine-coding stretches in the human genome (Butland et al., 2007) (Fig. S1A). Second, Hcf1, which is a pivotal binding partner of Ronin (Dejosez et al., 2008; Dejosez et al., 2010), was found to be part of the ataxia protein-protein interaction network (Lim et al, 2006), and has been broadly implicated in neurodevelopmental disorders including non-syndromic X-linked mental retardation (Ferrari et al., 2016; Huang et al., 2012; Vrabec et al., 2008; Turner et al., 2015; Jolly et al., 2015). Finally, mutations in Ronin’s highly similar gene cousin Thap1, causes DYT6 dystonia (Fuchs et al., 2009; Mazars et al., 2010), which is an inherited movement disorder that affects basal ganglia and the cerebellum (Neychev et al., 2011; Prudente et al., 2014; Nibbeling et al., 2017; Ruiz et al., 2015).

While evidence linking Ronin to SCA4 has been acknowledged (Pandey et al., 2004), sequencing efforts have failed to reveal any suspicious mutation or polyglutamine-coding repeat expansion in the SCA4-linked genomic region on chromosome 16q22.1 (Hellenbroich et al., 2005, Edener et al. 2011), and no abnormal polyglutamine aggregates were found in SCA4 patients (Hellenbroich et al., 2006). Hence, we argued, if Ronin were to be involved in the development of SCA4 or any other SCA, it would most likely entail alternative genetic causes. We hypothesized that the genetic alteration might involve aberrations such as an increased gene dosage (copy number variation, CNV) or non-coding changes that alter mRNA stability or modulate the expression of its encoding gene. To test this idea, we investigated whether ectopic expression of Ronin in Purkinje cells can induce ataxia in mice.

## RESULTS

### Ronin is present in Purkinje cells throughout the cytoplasm and nucleus

Spinocerebellar ataxias are caused at least in part by cerebellar Purkinje cell degeneration. Hence, we first asked if Ronin is expressed in Purkinje cells. In situ hybridization data (Allen Mouse Brain Atlas, Lein et al., 2007) and our previous expression analysis in Ronin-lacZ reporter mice (Dejosez et al., 2008) already suggested that *Ronin* is expressed in this cell type on RNA level. In line with this observation, immunostaining of sagittal cerebellar sections (Fig. 1A) confirmed that Ronin protein is prominently expressed in Purkinje cells. However, its subcellular localization differed from what we and others observed in other cell types, including embryonic stem cells (Dejosez et al., 2008) and tumor cells (Parker et al., 2012), where Ronin protein is primarily located in the nucleus. In Purkinje cells, Ronin protein is prevalent in the nucleus and the cytoplasm, suggesting that it is not only regulated by expression but also localization and/or that it might fulfill additional functions in the cytoplasm.

**Fig. 1.**
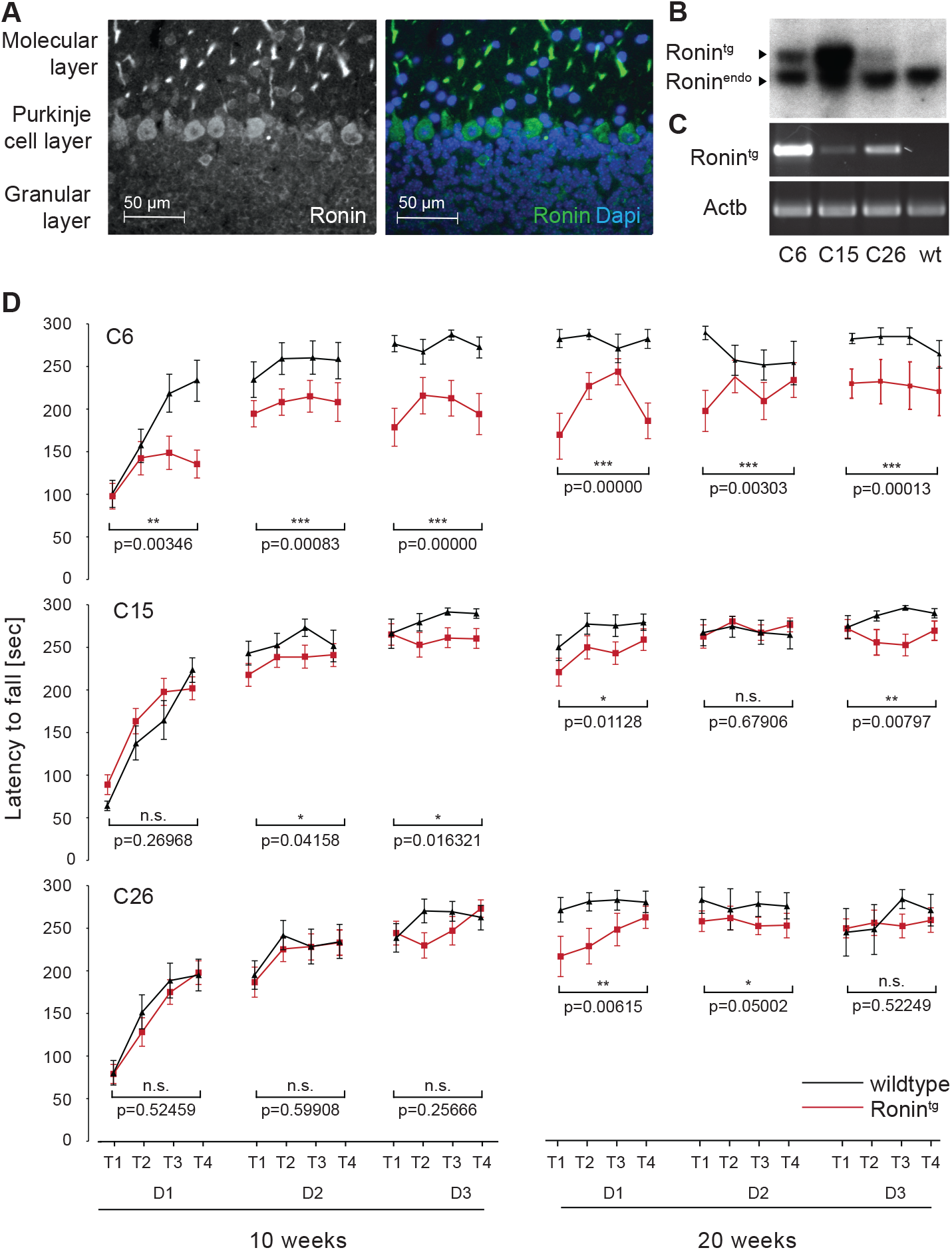
Ronin is expressed in Purkinje cells and Purkinje cell-specific transgenic expression of Ronin leads to reduced rotarod performance. (**A**) Immunofluorescence microscopy of 40 µm sagittal cerebellar paraffin sections of 12-week-old animals, stained with a Ronin specific antibody. Within the Purkinje cells, Ronin appears to be mainly located in the cytoplasm. (**B**) Southern blot analysis of transgenic animals in comparison to a wildtype (wt) control with a probe detecting the L7-*Ronin* transgene (Ronin^tg^) and endogenous *Ronin* (Ronin^endo^). (**C**) RT-PCR analysis of mRNA isolated from cerebella of transgenic and wildtype animals detected transgenic Ronin RNA in all three transgenic mouse lines. (**D**) Rotarod analysis of a representative set of animals from transgenic L7-Ronin (Ronin^tg^) lines (C6, C15 and C26) at 10 weeks (left) and 20 weeks (right) of age in comparison to wildtype littermates. Shown is the latency to fall in seconds in 4 trials (T1-T4) on 3 consecutive days (D1-D3). Transgenic animals show a reduced latency to fall. Data are shown as mean ± SEM; n=13, 24 or 13 for transgenic and n=14, 13 or 7 for wildtype animals of lines C6, C15 and C26 respectively.

### Purkinje cell-specific transgenic expression of Ronin causes ataxia

While some SCAs are caused by genes with expanded polyglutamine encoding repeats (Butland et al., 2007) no suspicious mutation was found in any gene residing in the SCA4-linked genomic region on chromosome 16q22.1 (Hellenbroich et al., 2005, Edener et al. 2011). With regards to Ronin this may not be surprising given that Ronin’s polyglutamine coding CAG repeat is interrupted by CAA triplets making replication-dependent slippage expansion unlikely (Fig. S1B). An alternative cause that has not been considered yet, is increased gene dosage (e.g. CNVs) of Ronin. We therefore tested if transgenic expression of normal Ronin protein might be sufficient to cause ataxia. To accomplish this goal, we used a Pcp2/L7-Ronin-Flag construct (Fig. S2A; Smeyne et al., 1995) to direct transgenic Ronin expression specifically to the Purkinje cells (Fig. S2B, Movie 1) of transgenic animals (Fig. S2B, Movie 1). Using Southern blot analyses (Fig. 1B) and RT-PCR (Fig. 1C), we confirmed transgene integration and expression in three independent mouse lines.

To determine if the Purkinje cell-specific expression of transgenic *Ronin* can induce ataxia, we tested the rotarod performance of our transgenic mouse lines at 10 - 12 weeks of age. The latency to fall was determined over the course of 5 minutes on an accelerating rotarod (3 - 30 rpm) (Watatse et al., 2002). At this time point, transgenic animals of two of our three L7-Ronin lines already showed significant decreases in rotarod performance when compared with wildtype littermates (Figs. 1D, right). When the experiments were repeated with the same groups of animals at 20 weeks of age, transgenic animals of all three lines showed significant differences in comparison to wildtype animals on the first trial day (Fig. 1D, left). The performance matched theirs from the last day of the previous round, indicating that they exhibited a training effect. Thus, the inability to stay on the rotarod did not appear to be caused by a general learning defect. Transgenic line C6 showed the strongest phenotype (Fig. 1D) and over time developed a marked ataxic gait that was seen in footprint pattern analyses (Fig. 2A, Movie 2). Additional phenotypic changes seen only in the transgenic animals included development of kyphosis (Fig. 2B) and abnormal clasping behavior (Fig. 2C).

**Fig. 2.**
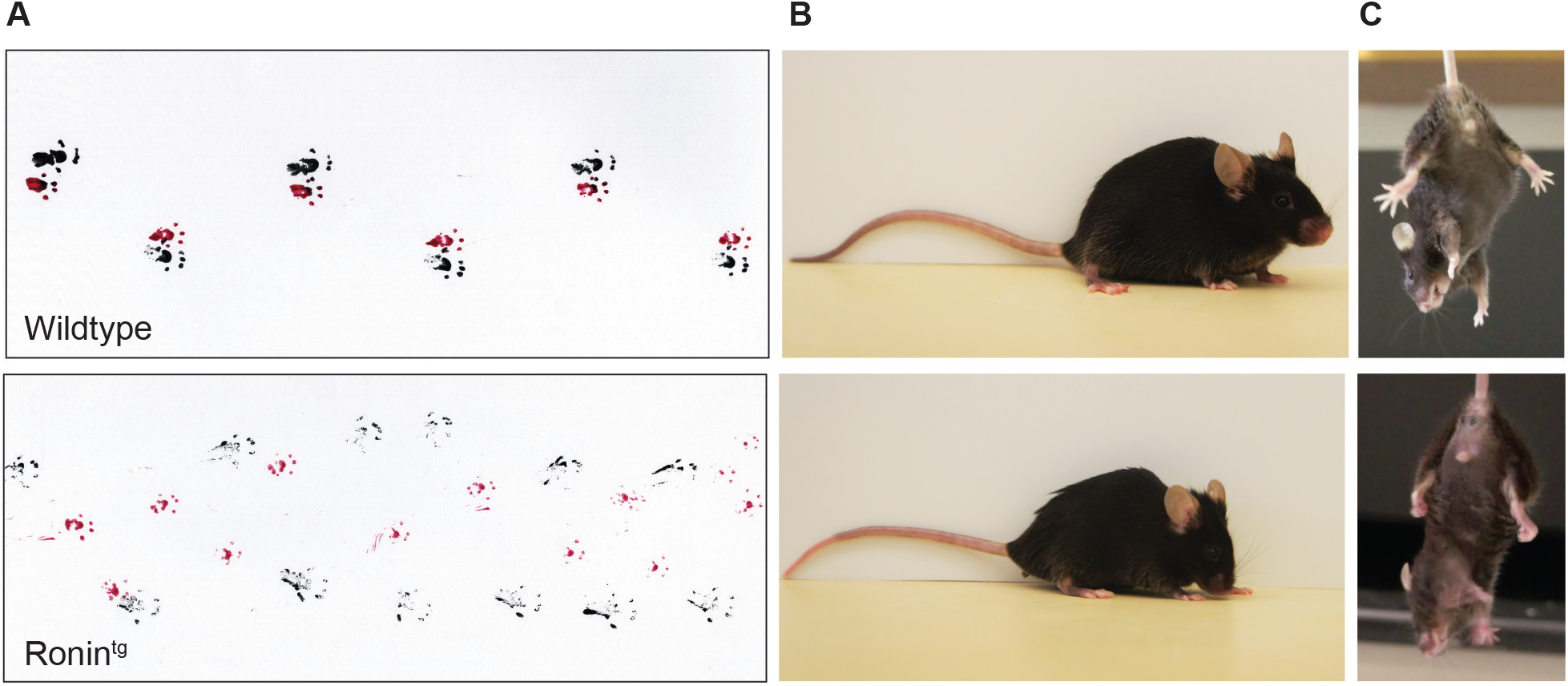
Purkinje cell-specific transgenic expression of *Ronin* leads to ataxia. Animals with Purkinje cell-specific transgenic expression of Ronin (Ronin^tg^) show signs of ataxia including an **(A)** abnormal gait pattern (red: front paws, black: hind paws), (**B**) a hunched back, and (**C**) abnormal clasping behavior when compared with wildtype litter mates. Animals shown are 1 year of age.

### Purkinje cell-specific transgenic expression of Ronin leads to cell loss

We next tested if the movement impairments and ataxic phenotype that we observed were related to Purkinje cell loss. At 5 weeks of age, transgenic cerebella were macroscopically indistinguishable from those of their wildtype littermates (Fig. 3A). Calbindin staining of floating cerebellar sections (Fig. 3B) and Hematoxylin-Eosin (HE) staining of paraffin sections (Fig. 3C) at this time point revealed similar numbers, morphology, and distribution of Purkinje cells in transgenic mice and wildtype littermates. In contrast, the same analyses at later time points of adolescence and adulthood (10, 20 and 58 weeks of age) revealed that there was a progressive decrease in Purkinje cell densities in transgenic animals compared to wildtype littermates (Figs. 4 and S4). Hence, we concluded that the neuronal loss was the result of neurodegeneration and not a developmental defect. By 1 year of age, the Purkinje cells were largely depleted in transgenic animals and their cerebella were substantially smaller than those of controls (Figs. 3C and S2B).

**Fig. 3.**
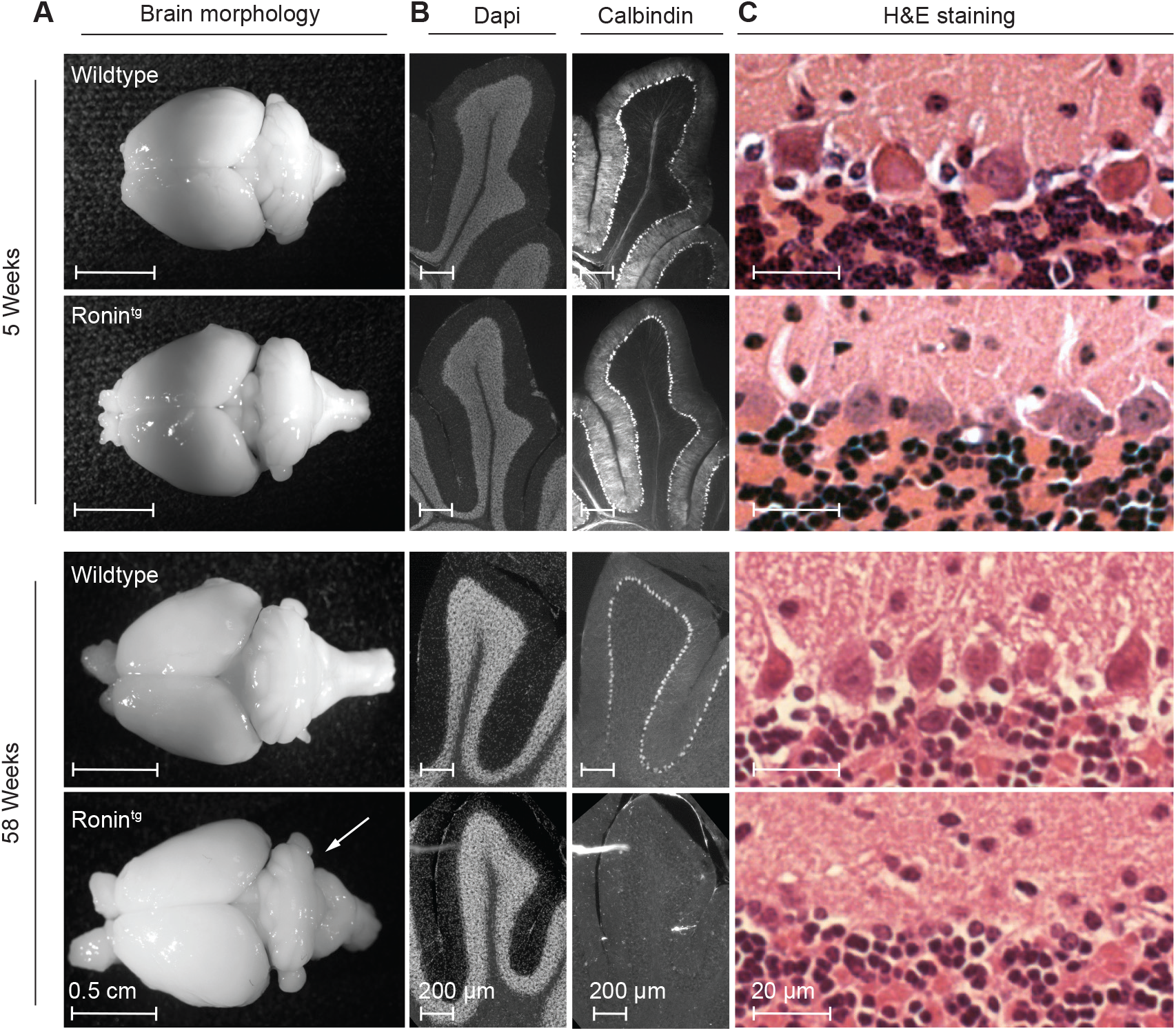
Purkinje cell-specific transgenic expression of *Ronin* leads to loss of Purkinje cells and cerebellar degeneration. (**A**) Macroscopic pictures of brains isolated from 58-week-old control or transgenic Ronin (Ronin^tg^) animals show drastically reduced size of the entire cerebellum when compared with wildtype controls (bottom), while at 5 weeks (top) there is no visible difference. **(B)** Immunofluorescence microscopy to detect Calbindin, a Purkinje cell-specific marker, and (**C**) Hematoxylin-Eosin (HE) staining of sagittal cerebellar sections demonstrates dramatic loss of Purkinje cells in Ronin^tg^ animals at 58 weeks of age, while at 5 weeks there is no discernable difference to wildtype controls.

**Fig. 4.**
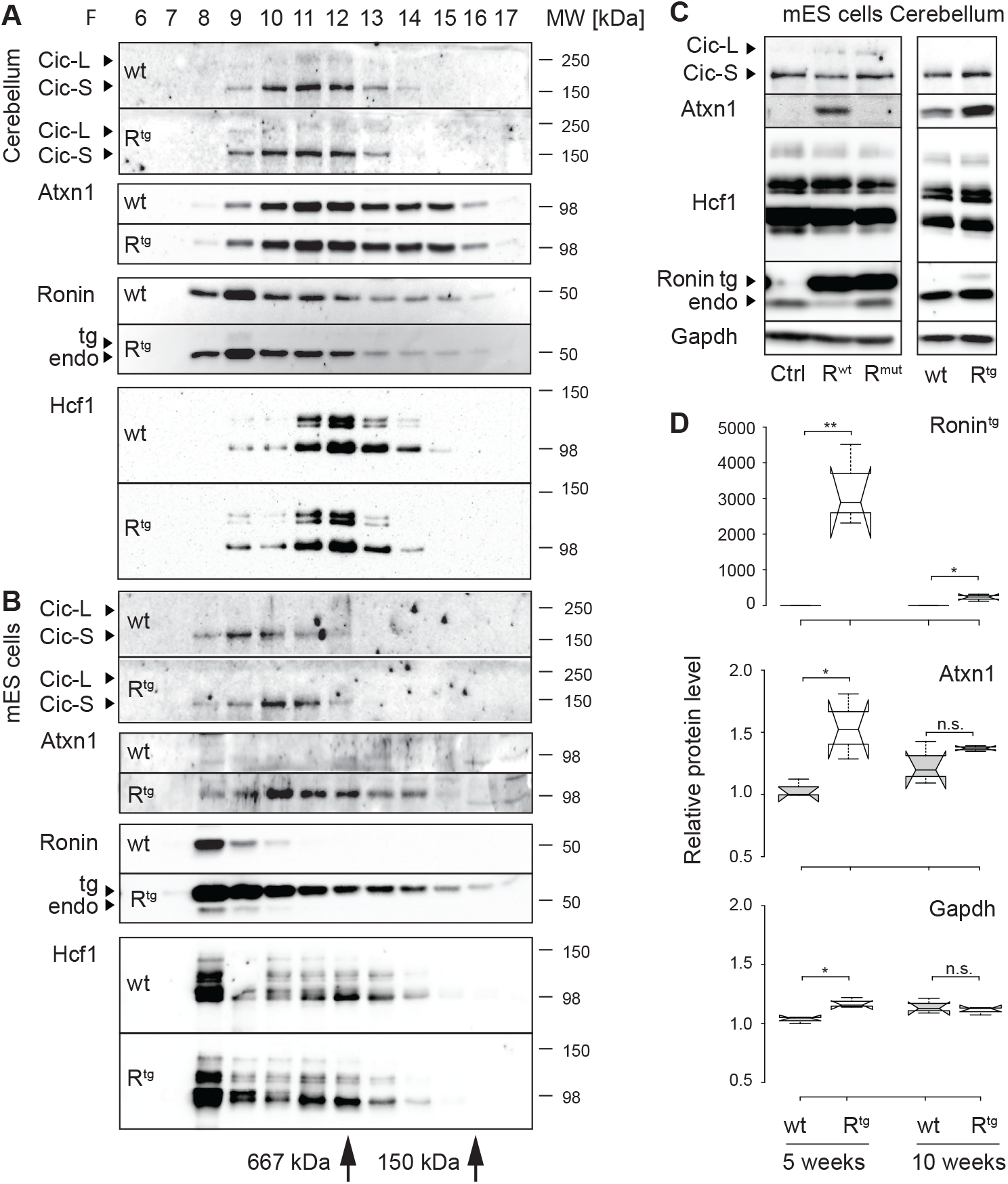
Transgenic expression of *Ronin* induces Atxn1. Representative Western blots of protein fractions collected by size exclusion chromatography of (**A**) cerebellar extracts or (**B**) mouse embryonic stem cell (mES) cell lysates. The size exclusion standards thyroglobulin (669 kDa) and ADH (150 kDa) are indicated. In the cerebellum, Ronin is part of a large protein complex that only partially overlaps with Hcf1. In ES cells, Atxn 1 expression is induced by transgenic expression of Ronin. Ronin and Hcf1 are eluting in similar fractions, indicating that they are part of the same protein complex. (**C**) Western blot analyses of equal protein amounts of ES cell lysates (left) or cerebellar extracts from 10-week-old animals (right) of indicated genotypes. While Atxn1 protein is not expressed in control ES cells overexpressing Luciferase (Ctrl), it is induced in Ronin overexpressing ES cells (R^wt^) but not in cells overexpressing a Ronin mutant incapable of binding to Hcf1 (R^mut^). Atxn1 levels are also elevated *in vivo* in transgenic (R^tg^) animals when compared to wt controls. (**D**) Quantification of transgenic Ronin, Atxn1 and Gapdh protein levels as detected by Western blot in cerebellar extracts at 5 and 10 weeks of age. At 5 weeks, a ~1.5-fold induction of Atxn1 is observed in transgenic Ronin animals when compared with wildtype littermates. Reflective of the decrease in Purkinje cells, at 10 weeks of age the expression of transgenic Ronin and Atxn1 induction were drastically reduced in comparison to the earlier time point. n=3; **p<0.01, *p<0.05.

### Purkinje cell-specific transgenic expression increases Ataxin-1 protein levels

We next addressed the question how, mechanistically, transgenic expression of Ronin might lead to Purkinje cell degradation and ataxia. In ES cells, Ronin’s function is dependent on its cofactor Hcf1, that had been identified as a component of the ataxia network described by Lim et. al. (2006). Within this network, the polyglutamine protein Ataxin-1 (Atxn1), the causative gene of the most common spinocerebellar ataxia SCA1, plays a central role. Atxn1 and Ronin are both factors that bind to DNA (Dejosez et al., 2010; Dejosez et al, 2008; Tsai et al., 2004) and form large protein complexes with common binding partners, such as SP1, Sin3A, Sap30 or the class 1 Histone deacetylase, HDAC3 (Weissman et al., 2018; Venkatraman et al., 2014; Dejosez et al., 2010). Furthermore, perturbation of the protein complex formed by Atxn1 and its binding partner Capicua (Cic), promotes the development of SCA1 (Lim et al., 2008, Rousseaux et al., 2019), thus providing a mechanistic basis for Purkinje cell loss. We therefore hypothesized that Ronin/Hcf1 and Atxn1/Cic are part of the same protein complex in the cerebellum, and that this complex might be disrupted in the cerebella of transgenic Ronin animals.

To test this prediction, we performed size-exclusion chromatography to analyze the distribution of Atxn1, Cic, Ronin and Hcf1 in cerebellar protein extracts in comparison with those of ES cells where Ronin is known to form a large protein complex (Dejosez et al., 2008) (Fig. 4A, B). We found that Hcf1 peaks partially overlapped with Atxn1 and Cic peaks, but there was only marginal overlap of Hcf1 with Ronin. The latter eluted with larger complexes and had a clear distinct peak in wildtype extracts. These results suggest that, Ronin and Hcf1 participate in two distinct protein complexes in the cerebellum. This is in contrast to the complexes found in ES cells (Fig. 4B), where Hcf1 peaked in the same fraction as Ronin, indicating that Ronin and Hcf1 are part of the same protein complex that we observed before (Dejosez et al., 2008; Dejosez et al., 2010). In addition, and different from Ronin, Hcf1 had a second elution peak in fractions similar to those in which it eluted in cerebellar extracts. Overall however, our data suggest that Ronin/Hcf1 and Atxn1/Cic do not cooperate in the same complex. In line with these results, we were unable to detect any interaction of Ronin with Atxn1 or Cic in immunoprecipitation studies using an anti-Flag antibody in ES cell extracts (Fig. S4A). Meanwhile, the previously described binding of Hcf1 to Ronin was readily visible (Dejosez et al., 2008).

While we did not find any differences in the distribution of any of the analyzed proteins between wildtype and transgenic samples per se, we noticed a dramatic increase in Atxn1 protein levels in EF1α-Ronin-Flag ES cells (Fig. 4B, C) when compared to controls. Indeed, Atxn1 was completely undetectable in control ES cells but emerged in Ronin overexpressing cells. This induction was dependent on Hcf1 since expression of a Ronin mutant incapable of binding to Hcf1 (Dejosez et al., 2010) did not show any Atxn1 protein (Fig. 4C, left) just like control cells. Therefore, Ronin is a potent inducer of Atxn1 *in vitro* and, importantly, Ronin also robustly induced Atxn1 protein levels in our transgenic *in vivo* model (Figs. 4C, S4B). Quantification of the protein levels (Fig. 4D) confirmed a ~1.5-fold induction of Atxn1 in cerebella of 5-week-old transgenic animals when compared to wildtype littermates. The induction was still detectable at 10 weeks of age but to a lesser extent, which can be attributed to the severe loss of Purkinje cells (Fig. 3) that is reflected in the substantial reduction of transgenic Ronin protein at this time point (Figs. 4D, S4B). The upregulation of Atxn1 is likely transcriptional in nature as Atxn1 expression was elevated on RNA level in Ronin overexpressing ES cells (Fig. S4C).

## DISCUSSION

We herein show that an increase in Ronin activity in cerebellar mouse Purkinje cells leads to Purkinje cell loss and development of severe ataxia in mice as early as 10-12 weeks after birth. Our data suggest that Ronin might be involved in the development of ataxia, including SCA4 and other neurodegenerative diseases, not through classical mutations in the gene body itself but rather through genetic defects that affect its activity (e.g., mutations in regulatory elements or copy number variations). At the mechanistic level, our data show that transgenic Ronin expression upregulates Atxn1, the causative gene of SCA1. This was an important finding, since recent studies have shown that an increase of wildtype Atxn1 protein leads to a gain-of-function of the Atxn1/Cic complex which is sufficient to induce ataxia in mice even in the absence of a polyQ expanded Atxn1 (Rousseaux et al., 2019). Because Ronin and its co-factor Hcf1, are associated with transcriptional regulation, the most parsimonious explanation for the increased Atxn1 levels in the presence of transgenic *Ronin* expression would be that Ronin exhibits transcriptional control over *Atxn1*. While *Atxn1* mRNA levels were indeed higher in Ronin overexpressing ES cells when compared to control cells (Fig. S4C; Dejosez et al., 2010), it remains to be determined, if Ronin affects Atxn1 through direct gene regulation or indirectly, such as by transcriptional regulation of genes involved in stabilizing or destabilizing *Atxn1* mRNA such as Pumilio (Gennarino et al., 2015). Pumilio is known to destabilize *Atxn1* mRNA, and its haploinsufficiency increases Atxn1 protein levels causing ataxia in mice (Gennarino et al., 2015) similar to the effects seen in our study.

Alternatively, Ronin may deregulate critical target genes through different molecular means altogether. For instance, its binding has been associated with 5-hydroxy-methylcytosines, and this finding links Ronin activity to the DNA methylation status at promoters (Spruijt et al., 2013). The DNA recognition sequence of Ronin is frequently found in chromatin loop anchors (Bailey et al., 2015), making it conceivable that some of its actions may be exerted through changes in the 3D genomic organization of promoters. Ronin’s gene regulation and DNA recognition sequence have been critically linked to heart development (Fujita et al., 2027), retina development (Poche et al., 2016) and disease (Radziwon et al., 2017) as well as cobalamin metabolism (Quintana et al., 2017). Additionally, polyglutamine proteins such as Ronin have recently been associated with the assembly of functional units (e.g., Pol II) through a biophysical process called phase separation (Toretsky and Wright, 2014). Given the DNA-binding ability of Ronin, its overexpression could alter gene expression by modulating Pol II recruitment. It is also noteworthy that Ronin was discovered as a caspase-3 target that has a functional caspase-3 cleavage site following the polyglutamine repeat (Fujita et al., 2017). Caspase cleavage products have been shown to contribute to the development of SCAs (Berke et al., 2004; Liman et al., 2014; Guyenet et al., 2015) and other neurological disorders (Graham et al., 2006). Thus, caspase-3 mediated cleavage might lead to accumulation of Ronin fragments which could contribute to the ataxic phenotype. Finally, Ronin was recently identified as a direct negative regulator of Parkin (PARK2), which is causing a familial form of Parkinson’s disease (autosomal recessive juvenile) when mutated (Potting et al., 2018). PARK2 is also known to interact with Ataxin 2 (ATXN2), whose polyglutamine expansion causes SCA2. Since PARK2 regulates the intracellular levels of both wildtype and mutant ATXN2 and inhibits ATXN2 induced cytotoxicity (Huynh et al., 2007), elevated expression of Ronin could lead to accumulation of ATXN2 through repression of PARK2.

It is remarkable that gene dosage changes of *Ronin* in its wildtype form can have such detrimental effects on neurons and induce such severe pathological outcomes. Given the robustness of the observed ataxia phenotype, it will be interesting to see if Ronin does, indeed, prove to be involved in 16q22.1-linked SCA4. In this case, we predict that genomic rearrangements (e.g., CNVs) might explain SCA4 disease and possibly other 16q22.1-linked ataxias. Such a scenario would add to the list of genomic rearrangement-related disorders, such as Charcot-Marie-Tooth disease (Parker et al., 2015), which is characterized by CNVs, SCA2, where the increased gene dosage in patients homozygous for the disease-related genetic defect influences the age of disease onset (Spadafora et al., 2007), autosomal dominant early-onset AD caused by a duplication of the amyloid precursor protein (APP) locus (Rovelet-Lecrux et al., 2006; Rumble et al., 1989), and familial Parkinson’s disease associated with duplications or triplications of α-synuclein (SNCA)(Chartier-Harlin et al., 2004; Ibanez et al., 2004; Singleton et al., 2003). Alternatively, simple mutations in *Ronin*’s regulatory elements (e.g., in one of its enhancers or the promoter) could cause chronic or context-specific misexpression to contribute to the disease. In light of *Ronin*’s genetically stable CAG repeat and the absence of polyQ conglomerates in SCA4 patients (Hellenbroich et al., 2006), our results suggest that such patients should be screened for duplications of the *Ronin* gene and possible mutations within the Ronin-binding motif of Ronin-bound promoters within the genomic region mapped to SCA4.

Future studies are needed to clarify whether expansion of the polyQ repeat in Ronin might lead to more severe outcomes. CAG repeat length differences have been described in both normal individuals and patients with neurological disorders (Yin et al., 2014) ranging from 22-38. 29 repeats are the most frequent (around 70%) followed by 28 repeats (~ 20%) with no significant difference between normal and diseased individuals. Interestingly, a 38Q repeat was detected in two neurodegenerative disease patients and expansion of the glutamine repeat *in vitro* leads to increased formation of nuclear aggregates in PC12 cells (Yin et al., 2014). Given the ataxic phenotype in our mouse model, we predict that further analyses of human Ronin, its binding partners, and its binding sites in target gene promoters might reveal novel mutations in the spectrum of neurodegenerative disease.

## MATERIAL AND METHODS

### Ronin immunofluorescence staining of paraffin sections

10-week-old animals were anesthetized and perfused with 4 % paraformaldehyde for 3 minutes. The brain was dissected and further fixed in 4 % paraformaldehyde overnight, transferred to 70°C and embedded in paraffin. 40 µM sections were stained with Ronin anti-serum G4276 (Dejosez et al., 2008) in a 1:1000 dilution for 16 hours at 4 °C using the TSA Kit 12 with HRP-goat anti Rabbit IgG and Alexa Fluor 488 tyramide conjugate (Invitrogen, T20922) following the manufacturer’s protocol. Stained sections were mounted in Vectashield with Dapi (Vector Laboratories).

### Generation of animals with Purkinje cell-specific transgenic expression of Ronin

To direct Ronin expression specifically to the Purkinje cells, a Not1/Sal1 fragment of pEF1α-hRonin-Flag-Ires-Neo (Dejosez et al., 2008), containing the human *Ronin* coding sequence, fused to a C-terminal Flag-tag, was cloned by blunt end ligation into the unique BamHI site of the pL7-AUG-EcoRI vector which is located in the fourth exon of the Pcp2/L7 gene (Smeyne et al., 1995). The vector includes approximately 1 kb of the promoter, the four exons and three introns as well as about 200 bp upstream sequence of the Pcp2/L7 gene as described. After sequence verification and HindIII digest, the construct was injected into single-cell blastocysts, that were transplanted into C57BL/6 foster animals by the Genetically Engineered Mouse Core at Baylor College of Medicine. All experimental procedures and protocols were approved by the Institutional Animal Care and Use Committee of Baylor College of Medicine (AN-4011) or the Icahn School of Medicine at Mount Sinai (IACUC-2013-1433), respectively. Founder animals were backcrossed to C57BL/6 animals to establish independent lines. L7-Ronin-Flag transgenic animals were identified by PCR of tail biopsies. DNA was isolated from 1-2 mm tail tissue using the DNeasy Blood and Tissue Kit (Qiagen) following the protocol for isolation of DNA from animal tissues and was eluted in 100 µl H_2_O. Subsequently, 2 µl DNA were used as template in two transgene-specific PCR assays. The PCR reaction was performed with the GoTaq Green Mastermix (Promega) in a total volume of 50 µl in the presence of 0.2 µM of each oligo. The L7-specific 5’ oligo MAD041 (TGT TTG GAG GCA CTT CTG ACT TGC) and the transgene specific oligo MAD224 (CTG ACT GCT GTC TAC AGT GGC CTG) amplified a 578 base pair fragment while the 5’ transgene specific oligo MAD225 (GCG GCC GCA AGA CCT ACA CGG TAC G) and the 3’ L7-specific oligo MAD042 (CAC TCA ACT CTT TGT TGC TAG TGC) resulted in a 969 base pair fragment. Following cycle conditions were used: denaturation for 3 min at 94 °C, 40 cycles of 30 sec at 94 °C, 30 sec at 55 °C and 30 sec at 72 °C, followed by a final extension at 72 °C for 7 min. Animals of both genders were included in this study. Controls were age- and gender matched if not otherwise stated.

### Southern blot analysis

To determine the copy number of the integrated transgene, Southern Blot analysis was performed as previously described with a *Ronin* specific probe complementary to part of the C terminus (Dejosez et al., 2008).

### RNA isolation and RT-PCR from cerebellar tissue samples

Cerebella were collected from 5-week-old animals and subjected to dounce homogenization. RNA from individual cerebella was isolated after homogenization with the Lipid Tissue RNA Kit (Qiagen) following the manufacturer’s instructions. 1 µg of RNA was used to create cDNA with the ImpromII Reverse Transcriptase Kit (Promega) following the manufactures standard protocol including oligodT oligos and 4.8 M MgCl2 in each 20 µl reaction (Promega). 1 µl of each reaction was used as template in subsequent PCR amplification reactions in a final volume of 50 µl with the GoTaq Green Mastermix (Promega) in the presence of 0.2 M of each oligo under following conditions: denaturation at 94 °C for 3 min, 30 cycles of 94 °C for 30 sec, 55 °C for 30 sec and 72 °C for 30 sec, followed by a final extension of 72 °C for 7 min. To detect the *hRonin*-Flag transgene mRNA the oligos MAD041 and MAD224 (see above) were used, while Actb mRNA was detected with the oligos MAD233 (GGC CCA GAG CAA GAG AGG TAT CC) and MAD234 (ACG CAC GAT TTC CCT CTC AGC) resulting in PCR fragments of 578 and 466 base pairs respectively.

### Rotarod analysis

Motor function was analyzed on a rotarod (UGO BASILE) accelerating from 4-40 rpm for 300 seconds and latency to fall was recorded in 4 consecutive trials on 3 consecutive days (Watase et al., 2002). All animals were from the second generation. 26 animals (11 tg/15 wt) of line C1, 27 animals of line C6 (13 tg/14 wt), 37 animals of line C15 (24 tg/13 wt) and 20 animals of line C26 (13 tg/7 wt) were tested. Animals in each group were born in a range of 2 weeks. Females and males were included in each group. P-values were calculated using two-tailed t-tests.

### Footprint analysis

Footprint analysis was done as previously described (Chen et al., 2012). Briefly, fore- and hind-paws of 50-week-old animals were painted with non-toxic red or black paint. The animals were allowed to walk through a tunnel lined with a sheet of paper.

### Calbindin staining of floating cerebella sections

Immunofluorescence staining of floating sections after perfusion was performed as described (Baader and Schilling, 1996). Briefly the animals were anesthetized and perfused with 4 % paraformaldehyde (PFA) for 3 minutes. The brain was dissected, further fixed in 4 % PFA for 16 hours and then treated in 20 ml of 5 %, 10 %, 15 %, 20 % and 25 % sucrose for at least 1 h or until the tissue sank to the bottom of the reaction tube. The brain tissue was cut in half and frozen in optimal cutting temperature (OCT) medium. 50 µM sections were cut and OCT was removed by two washes in PBS. Sections were blocked for one hour in blocking solution (2 % normal goat serum, 0.3 % Triton X-100 in DPBS), followed by incubation at 4°C in a 1:500 dilution of the anti-Calbindin-D-28K antibody (Sigma, CB-955) in blocking solution for 48 hours. Sections were washed 4 times for 20 minutes at room temperature and incubated with a 1:500 dilution of the goat anti-mouse Alexa-Fluor488 antibody (Molecular Probes) in blocking solution for 48 hours at 4 °C. The sections were washed 4 times at RT for 20 minutes, rinsed in PBS, transferred to microscope slides in 2 % BSA solution, dried and mounted in ProLong Gold with DAPI. Animals of both genders were analyzed and compared to age- and gender-matched controls.

### Hematoxylin-Eosin (HE) staining

Cerebella were fixed after perfusion, embedded in paraffin and 10 µm sections were stained with hematoxylin and eosin by the Histology Service of the Department of Pathology at Baylor College of Medicine.

### Cell culture

EF1α-Ires-Neo, EF1α-Ronin-Flag-Neo and EF1α-Ronin^DHSA^-Flag-Neo (Ronin^mut^) ES cells were cultured as previously described (Dejosez et al., 2010).

### Column fractionation

Complex purification and Western blot analyses were performed as previously described (Lam et al., 2006). Briefly, chromatography was carried out at 4°C using the GE AKTA Purifier UPC 10 FLPC system. Cellular or cerebellar extracts were prepared fresh by dounce-homogenization of ~25 million ES cells or two cerebella from age-matched mice (10-12 weeks) in TST buffer (50 mM Tris pH 8, 75 mM NaCl, 0.5 % Triton X-100, 1 mM PMSF, and protease inhibitors). 700 µl were loaded for FPLC. Gel filtration was a Superose 6 10/300 GL column (GE) equilibrated in buffer (50 mM Tris, pH 8, 50 mM NaCl, 0.1 % Triton X-100) at 0.3 ml/min. Fractions were collected every 0.5 ml or 1 ml volume, column void volume was 7.7 ml, and elution volumes of gel-filtration standards were 12.4 ml for thyroglobulin (669 kDa), 15.8 ml for ADH (150 kDa), and 19.2 ml for cytochrome C (12.4 kDa). Anion exchange was a MonoQ 5/5 HR column (GE) equilibrated in buffer (50 mM Tris, pH 8, 50 mM NaCl, 0.1 % Triton X-100); after washing the column with 13 column volumes of the same buffer, bound proteins were eluted with a 20 ml linear NaCl salt gradient from 50 mM to 600 mM NaCl in 50 mM Tris, pH 8, 0.1 % Triton X-100 at 1 ml/min; 1 ml fractions were collected. All fractions were supplemented with protease inhibitors and immediately prepared for SDS-PAGE.

### Protein analyses by Western blot

SDS page and Western blotting were performed using standard procedures (Chen et al., 2003, Watase et al., 2002). Following antibodies were used: guinea pig polyclonal anti-Cic serum (Chen et al., 2012), mouse monoclonal anti-FLAG M2 (F7425; Sigma), rabbit polyclonal anti-Atxn1 (11750VII) (Lam et al., 2006), mouse monoclonal anti-GAPDH (2-RGM2; Advanced Immunochemical), mouse monoclonal Ronin (Becton Dickinson), rabbit polyclonal Hcf1 (A301-400A, Bethyl Laboratories). Signal quantification was performed with the Image J software (Version 1.51, NIH). P-values were calculated using two-tailed t-tests.

## Acknowledgements

We thank Dr. Huda Y. Zoghbi for sharing her resources, her advice on experimental design, and critically reading the manuscript. We also thank Marco Passeri for performing Southern blots and cryosectioning, Marijke Schrock for the Ronin immunostaining, and Corinne Spencer from the Neurobehavioral Core at Baylor College of Medicine for assistance with the rotarod analysis.

## Author contributions statement

M.D. and T.P.Z. conceptualized the study, designed experiments, analyzed and interpreted data, and wrote the manuscript. M.D. and R.R. performed the experiments.

## Supplementary Information

Supplementary information consists of 4 supplementary figures (Figs. S1 to S4) and 2 movies (Movies 1 and 2).

